# Differential metabolic biomarkers between coronary heart disease and diabetes mellitus

**DOI:** 10.1101/2024.12.16.628812

**Authors:** Zhicai Wang, Yan Cang, Fei Shi, Yi Zhang, Hui-Na Cui, Yun He, Li Liu, Yan Li, Yawei Xu, He Wen, Zheng Liu

**Affiliations:** Department of Cardiology, Shanghai Tenth People’s Hospital, Tongji University School of Medicine, Shanghai, China; Pan-Vascular Research Institute, Heart, Lung, and Blood Center, Tongji University School of Medicine, Shanghai, China; Metanotitia Inc, Building C4, Science and Technology Innovation Headquarters, Shenzhen (Harbin) Industrial Park, 288 Zhigu Street, Songbei District, Harbin 150029, China; Cryo-electron Microscopy Center, Southern University of Science and Technology, Shenzhen, China

**Author notes:** Corresponding authors He Wen: E.mail Zheng Liu: E.mail. These authors contributed equally: Zhicai Wang and Yan Cang.

**Keywords:** Metabolomics, Coronary heart disease, diabetes mellitus, disease-specific biomarkers

## Abstract

**Background:** Coronary heart disease (CHD), the leading cause of death globally, is a complex disease often association with metabolic disorders, particularly when co-occurring with diabetes mellitus (DM). However, the mechanisms underlying the interaction between these two chronic conditions remain poorly understood.

**Methods:** In this study, we enrolled 320 participants, including 103 healthy controls (HC), 62 individuals with CHD, 44 with DM, and 47 with both CHD and DM (CHDDM)) in the training set, along with 64 participants (20 HC, 18 CHD, 12 DM, and 14 CHDDM) in the test set. Plasma metabolomic profiling was performed using gas chromatograph-mass spectrometry (GC-MS) and ultra-high performance liquid chromatography-mass spectrometry (UHPLC-MS)-based multi-platform untargeted metabolomics.

**Results:** Using multi-platform metabolomics, we identified 453 distinct metabolites. Through multivariate statistical analysis, we assigned 63, 27, and 56 disease-associated metabolic biomarkers for CHD, DM, and CHDDM, respectively, followed by pathway enrichment analysis. Using an area under the receiver operating characteristic (ROC) curve (AUC) cutoff value > 0.77, we identified 10, 4, and 8 disease-specific biomarkers for CHD, DM, and CHDDM, respectively. Notably, N-Formyl-L-methionine, N-Acetyl-L-methionine, PE 36:1p (18:0p/18:1), PE 36:2p (18:0p/18:2), and PE 34:2p (16:0p/18:2) were predominant in CHD, glycerophosphocholine and turanose were distinctive to DM, and xanthine and PE 38:5p (18:1p/20:4) were unique to CHDDM. These disease-specific metabolites showed superior diagnostic performance in their respective conditions. Additionally, 23 metabolites were identified as potential biomarkers for assessing the risk of CHD in DM patients.

**Conclusion:** This comparative metabolomics approach provided new insights into disease-specific markers and pathogenic pathways, offering potential for improved diagnosis and management of CHD and DM.

## Introduction

Coronary heart disease (CHD) is the leading cause of death globally, recognized as a multifactorial disease influenced by a range of complex metabolic disorders (1). Recent clinical trials and epidemiological studies have significantly enhanced our understanding of CHD pathogenesis, particularly its strong association with conditions such as hypertension, hyperlipidemia and diabetes mellitus (DM)(2) Among these, DM stands out as a particularly significant risk factor, individuals with diabetes have a 2- to 4-fold higher risk of developing CHD compared to non-diabetic individuals (3).

DM is not only a major independent risk factor for cardiovascular disease like CHD, but it also exacerbates the mechanisms involved in atherosclerosis, endothelial dysfunction, and prethrombotic status (4). A pivotal study questioned whether diabetes should be regarded as a CHD-equivalent condition, finding that the lifetime risk of vascular death in individuals with DM but no prior CHD was comparable to those with CHD alone (5). Beyond increasing the risk of CHD, DM significantly worsens the prognosis and mortality rates among CHD patients (6). Research indicates that approximately one-third of CHD patients are also diagnosed with DM (CHDDM), and these individuals face a poorer long-term prognosis (7). This could be due to the higher disease burden and more complex lesions in CHD patients with concurrent diabetes. However, the specific metabolic biomarkers and pathways involved in CHD and/or DM remain inadequately understood, underscoring the urgent need to elucidate the metabolic regulatory mechanisms in these diseases.

CHD and DM, as prevalent chronic diseases, are linked by intricate metabolic disorders, many of which remain to be fully explored. Metabolomics, a powerful approach for profiling small molecule metabolites, has increasingly been utilized in CHD research (8). Biomarker analysis, in particular, shows great promise for the early diagnosis and risk prediction of DM or CHD. It is well-established that significant metabolic alterations are integral to the onset and progression of CHD, with amino acids playing a critical role as key nutrients for cell growth and survival (9). Evidence suggests that defects in amino acid catabolism, particularly in branched-chain amino acids (BCAAs) like isoleucine, leucine, and valine, can exacerbate adverse myocardial remodeling and cardiac dysfunction following myocardial infarction by activating mammalian target protein (mTOR) signaling(10). While the regulatory role of amino acids in DM has been extensively studied, for instance, increased levels of BCAAs and aromatic amino acids (phenylalanine, tryptophan, and tyrosine) are associated with DM (11), suggesting the distinct but related biological mechanism between CHD and DM warrant further investigation.

Given the intertwined nature of CHD and DM, employing metabolomics to identify biomarkers and key metabolic pathways in patients with CHD and/or DM can provide deeper insights into their pathophysiological connections. Such insights are crucial for developing effective early diagnostic strategies, which are essential for the clinical management of CHD and CHDDM.

In this study, we investigated plasma metabolomic profiling using a combination of gas chromatography-mass spectrometry (GC-MS) and ultra-high performance liquid chromatography-mass spectrometry (UHPLC-MS). Through multivariate statistical analysis, we identified disease-associated metabolites and conducted pathway enrichment analysis. Univariate receiver operating characteristic (ROC) analyses, adjusted for variables, revealed 10, 4, and 8 specific biomarkers for CHD, DM, and CHDDM, respectively, with several biomarkers demonstrating superior diagnostic performance in their respective conditions. Additionally, 23 metabolites were identified as potential biomarkers for assessing CHD risk in DM patients. This comparative metabolomics approach offers valuable insights into disease-specific metabolic biomarkers and the underlying metabolic pathways, paving the way for improved diagnostic and therapeutic strategies.

## Materials and Methods

### Chemicals and reagents

Liquid chromatography (LC) and mass spectrometer (MS)-grade formic acid, acetic acid, ammonium acetate, methanol, acetonitrile, isopropanol, and water were commercially purchased from Sigma-Aldrich (St Louis, MO, USA).

### Study subjects

A total of 256 participants, including 103 healthy controls (HC), 62 patients with coronary heart disease (CHD) alone, 44 patients with diabetes mellitus (DM) alone, and 47 patients with both CHD and DM (CHDDM), were recruited as training set to define biomarker candidates. In order to validate biomarker candidates, blood samples from 20 healthy controls (HC), 18 patients with CHD alone, 12 patients with DM alone, and 14 patients with both CHD and DM (CHDDM) were collected as test set. The CHD participant inclusion criteria include symptoms of chest pain, ischemic changes in ECG, cardiovascular risk factors, or elevated myocardial enzymes. All the CHD patients were diagnosed and confirmed through coronary angiography. Coronary angiography shows that any coronary artery stenosis is greater than or equal to 50%, and is diagnosed as coronary heart disease by two or more experts, The patients with DM were diagnosed according to standard clinical criteria (12). The clinical diagnostic criteria for diabetes are as follows: fasting plasma glucose (FPG)≥7.0 mmol/L or 2-h plasma glucose (2-h PG) ≥11.1 mmol/L during oral glucose tolerance test (OGTT). The diagnosis of CHDDM is established by integrating the aforementioned criteria for both DM and CHD. The study recruited participants aged 18-65 years old who were not pregnant and did not have other chronic conditions, such as hypertension and chronic bronchitis. Patients were excluded if they had aortic dissection, pulmonary embolism, pneumonia, pericarditis, myocarditis, stress-induced cardiomyopathy, severe heart failure, liver or kidney failure, and blood-borne infections such as HIV/AIDS, hepatitis B, or hepatitis C. Additionally, patients with a history of malignant tumors, autoimmune diseases, severe traumatic diseases, infectious diseases, or who had undergone traumatic surgery within the last 3 months were also excluded. This study was evaluated by the Research Ethics Committee of the Tenth People’s Hospital of Shanghai, No. 20K149 and 21K65. All participants agreed and signed informed consent forms. The study was conducted in accordance with the guidelines of the International Conference on Harmonization for Good Clinical Practice and the Declaration of Helsinki.

### Sample preparation

Blood samples were obtained from all participants in fasting condition. One milliliter blood was collected in BD Vacutainer Serum Plus Tube for biochemical analysis. Five milliliter blood were obtained in EDTA-K2 anticoagulant tube and centrifuged at 3,800 rpm at 4℃ for 12 min. The plasma samples in supernatant were moved in EP tube and stored at −80℃ until analysis.

Fifty microliter plasma sample mixed with 700 μL of extraction buffer containing methanol/methyl tert-butyl ether/water (v/v/v, 1:3:1) containing 0.45 μg/mL of gibberellic acid A3, 1 μg/mL of ^13^C sorbitol, and 0.45 μg/mL of PE (17:0/17:0) as internal standards. Gibberellic acid A3 and PE (17:0/17:0) were used as the internal standard of hydrophilic and lipophilic phase detection LC-MS platform, respectively. In addition, ^13^C sorbitol was applied for GC-MS platform. Internal standards were utilized to monitor the stability of extraction and the on-board process of the MS platform. The coefficient of variation (CV) for each substance less than or equal to 20% was considered as stable.

Then, the samples were sonicated for 15 minutes in a 4 °C bath. Subsequently, 350 µL solution (methanol/water, v/v, 1:3) was added to facilitate phase separation. After The upper lipophilic phases and lower hydrophilic phases were separated by high-speed centrifugation (12,700 rpm, 5 min at 4 °C, Centrifuge 5430R, Eppendorf, Germany). Next, 400 µL of the lower hydrophilic phase was further mixed with 1.1 mL of methanol, incubated at 4 °C for 1 hour, and then centrifuged at 12,700 rpm for 10 minutes at 4 °C. The upper phase containing the lipophilic phase (350 µL) was transferred to a microcentrifuge tube for lipid analysis. The hydrophilic phase was divided into two microcentrifuge tubes, one containing 350 µL and the other containing 1,000 µL, intended for GC-MS and UHPLC-MS metabolite analysis, respectively. All aliquots were dried using a speed vacuum concentrator and stored at −80 °C. Before use, the dried samples of lipophilic and hydrophilic constituents were dissolved in 200 μL of ACN/IPA (v/v, 7:3) and water, respectively. Finally, they were transferred into sample vials for further analysis.

To ensure measurement quality and equipment performance, three types of quality control (QC) samples were prepared according to the aforementioned procedure. These included a pooled sample consisting of 50% randomly selected DBS samples (QCbio), a mixture of chemical standards (QCmix), and a sample containing only solvents (QCblank). To identify impurities in the solvents or contamination in the separation system, the analytical batch run began with a QCblank. QCmix and QCbio samples were inserted after every ten biological samples during analysis.

### GC-MS and UHPLC-MS Detection

The derivatization of the dried hydrophilic metabolites was performed according to Lisec et al. (13). Briefly, the dried fractions were oximilised by methoxyammonium pyridine solution and derivatized by adding MSTFA for GC-MS analysis. The analysis of derivatization of the hydrophilic metabolites was conducted utilizing an Agilent 7890B gas chromatograph with an Rxi®-5SilMS GC column (30 m, 0.25 × 30 mm, 0.25 µm), coupled to time-of-flight (TOF) mass spectrometer (Leco Corp., St. Joseph, MI, USA) with an electron ionization source by an Agilent 7683 series autosampler (Agilent Technologies GmbH, Waldbronn, Germany). Data were acquired both in positive and negative modes. High-purity helium was set as carrier gas with a flow rate of 1 mL/min. The temperature was initially set at 50 °C for 2 min, and then reaching 330 °C at a rate of 1 °C per minute. The ion source temperature and interface were set at 250 °C and 280 °C, respectively. The detector voltage and electron energy were set at 1.2KV and 70eV, respectively.

The analysis of lipophilic and hydrophilic extractions was performed using an ultra-high performance liquid chromatography (UHPLC, Waters ACQUITY (Milford, MA, USA)) system coupled to Thermo-Fisher Q-Exactive (Bremen, Germany) mass spectrometers with an electrospray ionization (ESI) source under both positive and negative modes. For lipophilic samples, a BEH C8 column (1.7µm, 2.1×100mm) was used for chromatographic separation with a column temperature of 60 °C. Gradient elution was performed using a mobile phase A water and B ACN/IPA (70/30, V/V), both containing 0.1% acetic acid and 0.1 % ammonium acetate at a flow rate of 0.4 mL/min. The optimal gradient elution program was set as follows: 55% B at 0-1 min, 75% B at 4 min, 89% B at 12 min, 100% B at 12 min, 100% B at 19.5 min, 55% B at 19.51 min, 55% B at 24 min. The injection volume of the sample was set at 2 μL with a temperature held at 10 °C. For hydrophilic samples, chromatographic separations were performed on a HSS T3 column (1.8µm, 2.1×100mm) with a column temperature of 40 °C. Gradient elution was carried out using mobile phase A water and B ACN, both containing 0.1% formic acid at a flow rate of 0.4 mL/min. The gradient elution program was operated as follows: 1% B at 0 min, 70% B at 13 min, 99% B at 13.01 min, 99% B at 18 min, 1% B at 18.01 min, and 1% B at 22 min. The injection volume of the sample was set at 3 μL with the temperature held at 10 °C. For both lipophilic and hydrophilic metabolite analyses, full scan and data-dependent acquisition (DDA) were employed to acquire MS and MS/MS spectra. Full scanning analyses were performed in the range of m/z 100–1,500 Da. The DDA was carried out for targeting the top five most intense precursor ions for MS/MS analysis in the pooled DBS samples (QCbio) with the scan ranges of m/z 100−310 Da, 300−710 Da, and 700−,1500 Da, respectively. The operating conditions of the mass spectrometer were as follows: 3500 V in positive mode and 3000 V in negative mode of ion spray voltage, 20 psi nebulizer, 400 sheath gas temperature, 10 L/min sheath gas flow, and 30 V normalized collision energy was used for MS/MS (14).

### Data processing and analysis

Chromatograms obtained from GC-MS analysis were exported by Leco ChromaTOF (version 5.40). Peak detection, fatty acid methyl esters (FAME)-based retention time alignment, and mass spectral comparison with the Fiehn reference libraries (15) were carried out using the Bioconductor package TargetSearch (16) in R (version 4.0.3). Metabolites were quantified by the peak intensity of a selective mass. Retention time (RT) deviation in GC-MS was calculated as the RT shift throughout all batches for each FAME internal standard (14).

UHPLC-MS chromatograms were processed using Metanotitia Inc. in-house developed software *PAppLine*^TM^. supplemented by commercial software including Compound Discoverer 3.1 and LipidSearch (Thermo Fisher Scientific, Waltham, MA, USA). The first processing step including peak detection, peak filtering, baseline correction and removing isotopic peaks(17). The lipids and metabolites were annotated based on the retention time, precursor ion and product ions fragmentation pattern by using the Metanotitia Inc. library, Ulib^MS^. In detail, the establishment of metabolite database via a sub-library of six thousand compounds, which were developed according to same chromatographic and spectrometric conditions as the measured samples, and the lipids were annotated with a sub-library of 1,700 lipids based on the precursor m/z, fragmentation spectrum, and elution patterns(14).The matching criteria were retention time within of 0.2 min and mass accuracy lower than 10 ppm. Then, the features that were detected in less than 50 % of the plasma samples were removed for further analysis. The missing values were imputed by using Mice random forest algorithm according to the sample type and peak intensity (17).Finally, a calibration procedure including logarithm transformation and scaling was performed. To reduce the difference of metabolite concentration among samples and make the data distribution more symmetrical, the Normalization Autoencoder (NormAE) was used(18).

### Statistical analysis

Multivariate statistical analysis was performed by using MetaboAnalyst 6.0. (https://www.metaboanalyst.ca/). Orthogonal partial least-squares discriminant analysis (OPLS-DA) was also used to identify the differences between groups. The statistical analysis and OPLSDA-based variable importance in the projects (VIP) analysis were also performed to investigate the differences of metabolites. Additionally, pathway enrichment analysis was also employed to elucidate the disease-related pathways. The Venn diagrams were generated using OmicStudio tools (https://www.omicstudio.cn/tool). Receiver-Operating Characteristic (ROC) Analysis was used to carry out diagnostic tests for the two groups with multiple compound combinations, and the area under the working curve (AUC) of the subjects served to assess the classification ability of the compounds involved in the two groups.

## Results

### Characteristics of the study subjects

Baseline characteristics of the study participants are summarized in **Table 1a** and **1b**. The training set included 256 participants: 62 with CHD, 44 with DM, 47 with CHDDM, and 103 healthy controls (HC). The test set comprised 62 participants: 16 with CHD, 12 with DM, 14 with CHDDM, and 20 HC. Across both sets, clinical and biological characteristics such as age, sex, blood urea nitrogen (BUN), low-density lipoprotein cholesterol (LDL-C), and high-density lipoprotein cholesterol (HDL-C) were similar among groups. Fasting glucose levels were significantly elevated in the DM and CHDDM group compared to the HC in both sets. In training set, body mass index (BMI) and triglycerides (TG) were lower in the HC group compared to all patient groups. In contrast, in test set, BMI was higher only in CHD and CHDDM group. Glycated hemoglobin (HbA1c) levels were significantly higher in all patient groups compared to the HC subjects both in training and test sets.

**Table 1.**
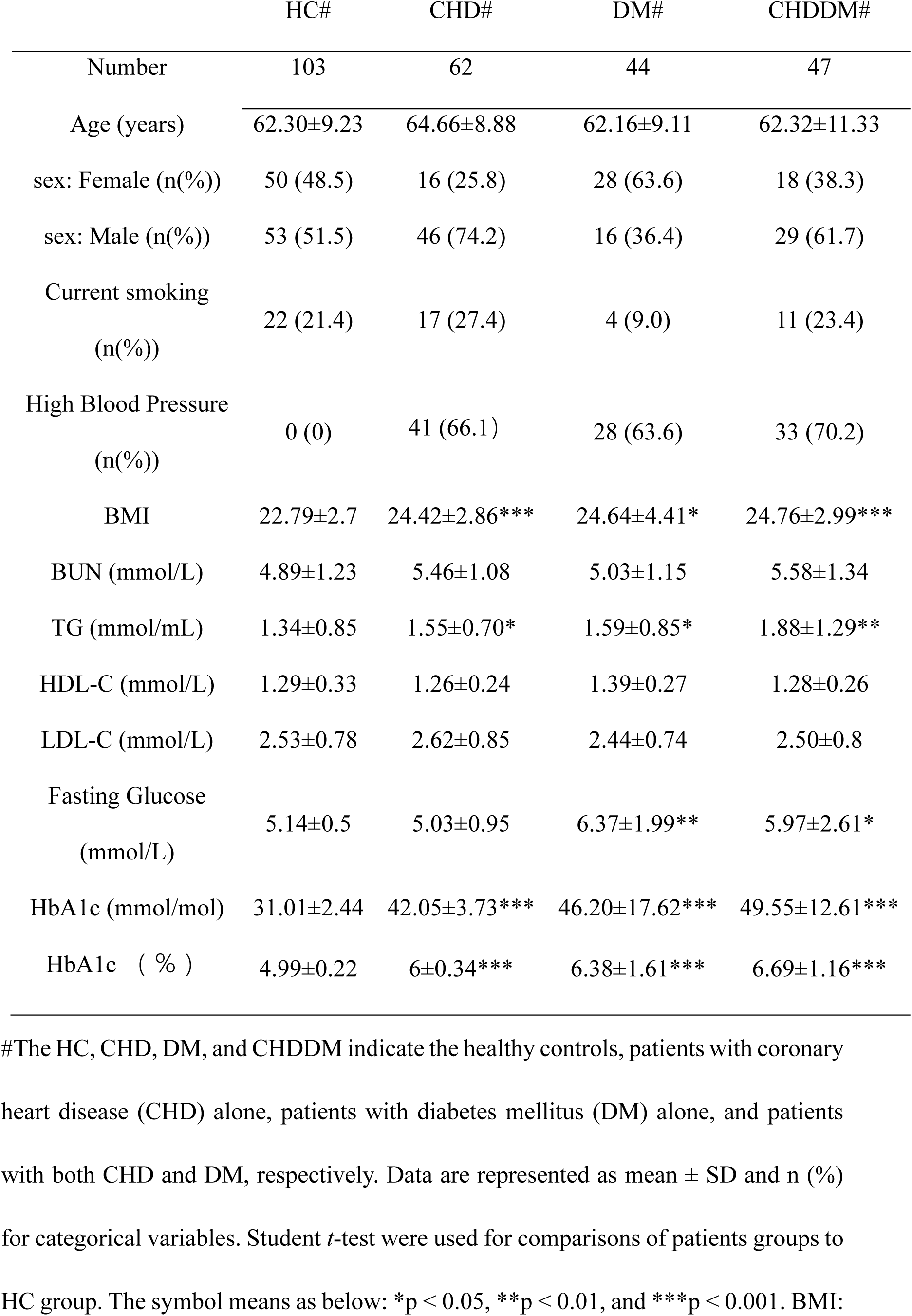
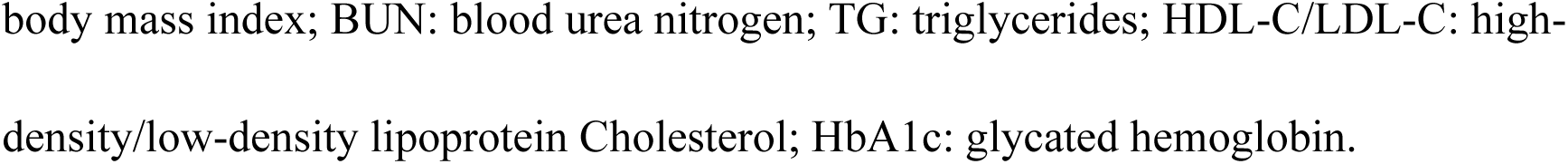
The basic information of the patients and healthy controls in training set.

|  | HC# | CHD# | DM# | CHDDM# |
| --- | --- | --- | --- | --- |
| Number | 103 | 62 | 44 | 47 |
| Age (years) | 62.30±9.23 | 64.66±8.88 | 62.16±9.11 | 62.32±11.33 |
| sex: Female (n(%)) | 50 (48.5) | 16 (25.8) | 28 (63.6) | 18 (38.3) |
| sex: Male (n(%)) | 53 (51.5) | 46 (74.2) | 16 (36.4) | 29 (61.7) |
| Current smoking<br>(n(%)) | 22 (21.4) | 17 (27.4) | 4 (9.0) | 11 (23.4) |
| High Blood Pressure<br>(n(%)) | 0 (0) | 41 (66.1) | 28 (63.6) | 33 (70.2) |
| BMI | 22.79±2.7 | 24.42±2.86*** | 24.64±4.41* | 24.76±2.99*** |
| BUN (mmol/L) | 4.89±1.23 | 5.46±1.08 | 5.03±1.15 | 5.58±1.34 |
| TG (mmol/mL) | 1.34±0.85 | 1.55±0.70* | 1.59±0.85* | 1.88±1.29** |
| HDL-C (mmol/L) | 1.29±0.33 | 1.26±0.24 | 1.39±0.27 | 1.28±0.26 |
| LDL-C (mmol/L) | 2.53±0.78 | 2.62±0.85 | 2.44±0.74 | 2.50±0.8 |
| Fasting Glucose<br>(mmol/L) | 5.14±0.5 | 5.03±0.95 | 6.37±1.99** | 5.97±2.61* |
| HbA1c (mmol/mol) | 31.01±2.44 | 42.05±3.73*** | 46.20±17.62*** | 49.55±12.61*** |
| HbA1c ( % ) | 4.99±0.22 | 6±0.34*** | 6.38±1.61*** | 6.69±1.16*** |

#The HC, CHD, DM, and CHDDM indicate the healthy controls, patients with coronary heart disease (CHD) alone, patients with diabetes mellitus (DM) alone, and patients with both CHD and DM, respectively. Data are represented as mean ± SD and n (%) for categorical variables. Student *t*-test were used for comparisons of patients groups to HC group. The symbol means as below: \**p* < 0.05, \*\**p* < 0.01, and \*\*\**p* < 0.001. BMI:

**Table 1b.**
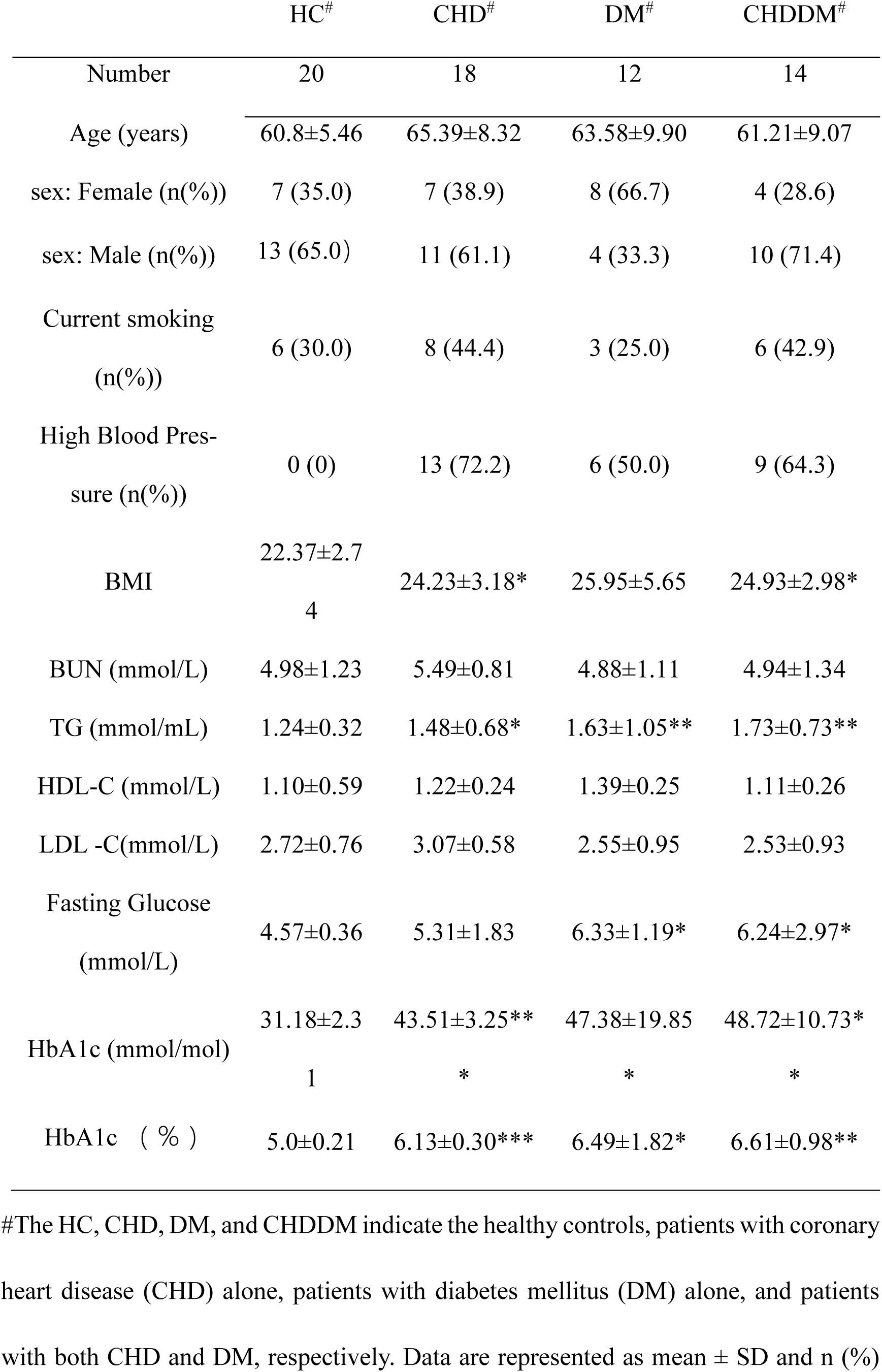
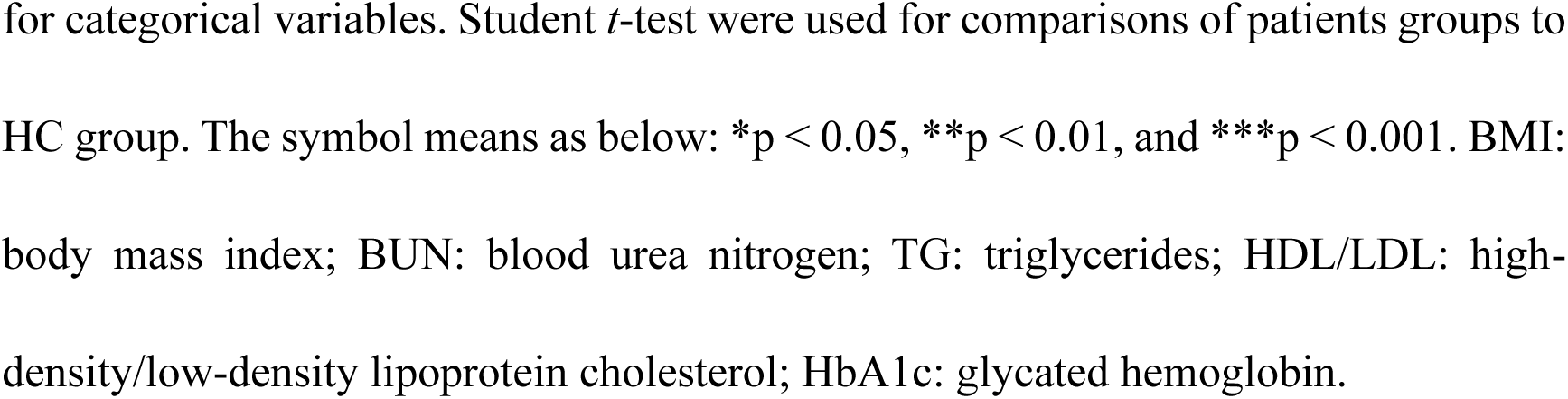
The basic information of the patients and healthy controls in test set.

|  | HC<sup>#</sup> | CHD<sup>#</sup> | DM<sup>#</sup> | CHDDM<sup>#</sup> |
| --- | --- | --- | --- | --- |
| Number | 20 | 18 | 12 | 14 |
| Age (years) | 60.8±5.46 | 65.39±8.32 | 63.58±9.90 | 61.21±9.07 |
| sex: Female (n(%)) | 7 (35.0) | 7 (38.9) | 8 (66.7) | 4 (28.6) |
| sex: Male (n(%)) | 13 (65.0) | 11 (61.1) | 4 (33.3) | 10 (71.4) |
| Current smoking<br>(n(%)) | 6 (30.0) | 8 (44.4) | 3 (25.0) | 6 (42.9) |
| High Blood Pres-<br>sure (n(%)) | 0 (0) | 13 (72.2) | 6 (50.0) | 9 (64.3) |
| BMI | 22.37±2.7<br>4 | 24.23±3.18* | 25.95±5.65 | 24.93±2.98* |
| BUN (mmol/L) | 4.98±1.23 | 5.49±0.81 | 4.88±1.11 | 4.94±1.34 |
| TG (mmol/mL) | 1.24±0.32 | 1.48±0.68* | 1.63±1.05** | 1.73±0.73** |
| HDL-C (mmol/L) | 1.10±0.59 | 1.22±0.24 | 1.39±0.25 | 1.11±0.26 |
| LDL -C(mmol/L) | 2.72±0.76 | 3.07±0.58 | 2.55±0.95 | 2.53±0.93 |
| Fasting Glucose<br>(mmol/L) | 4.57±0.36 | 5.31±1.83 | 6.33±1.19* | 6.24±2.97* |
| HbA1c (mmol/mol) | 31.18±2.3<br>1 | 43.51±3.25**<br>* | 47.38±19.85<br>* | 48.72±10.73*<br>* |
| HbA1c ( % ) | 5.0±0.21 | 6.13±0.30*** | 6.49±1.82* | 6.61±0.98** |

<sup>#</sup>The HC, CHD, DM, and CHDDM indicate the healthy controls, patients with coronary heart disease (CHD) alone, patients with diabetes mellitus (DM) alone, and patients with both CHD and DM, respectively. Data are represented as mean ± SD and n (%)
for categorical variables. Student *t*-test were used for comparisons of patients groups to HC group. The symbol means as below: \* $p < 0.05$ , \*\* $p < 0.01$ , and \*\*\* $p < 0.001$ . BMI: body mass index; BUN: blood urea nitrogen; TG: triglycerides; HDL/LDL: high-density/low-density lipoprotein cholesterol; HbA1c: glycated hemoglobin.

### GC-MS and UHPLC-MS detection and multivariate statistical analysis

To explore metabolic differences among disease groups and HC, plasma metabolic profiling was conducted using both positive and negative modes GC-MS and UHPLC-MS-based multi-platform untargeted metabolomics. A total of 453 distinct metabolites were identified, classified into various subclasses, such as amino acids, carbohydrates, fatty acids, glycerolipids (GL), glycorophopholipids (GP), nucleotides, organic acids, peptides, sphingolipids (SP), TCA cycle, steroids, and others (**Table S1**). A heatmap of the metabolomic data revealed clear differentiation between disease groups and HC group (**Figure S1**).

To evaluate the statistical significance of the metabolites and exclude potential confounding variables, orthogonal projections to latent structure-discrimination analysis (OPLS-DA) multivariate analysis was employed. The OPLS-DA distinction models were obtained with one predictive and at least two orthogonal components (see **Table S2**). The OPLS-DA models effectively distinguished the HC group from DM and CHDDM groups (**Figure 1b and** **1c**) and showed some separation between CHD and HC groups with slight overlap (**Figure 1a**). The discrimination model demonstrated robust goodness of fit (R^2^Y) and cross-validation (Q^2^) coefficients, with overall R^2^X values exceeding 0.767, 0.637, and 0.293, respectively (**Table S2**). We performed one thousand permutations, revealing that the resulting of R^2^Y and Q^2^ values were lower than the original values (**Table S2**). Permutation testing confirmed the validity of the OPLS-DA models, indicating high cross-validated predictability and goodness-of-fit.

**Figure 1.**
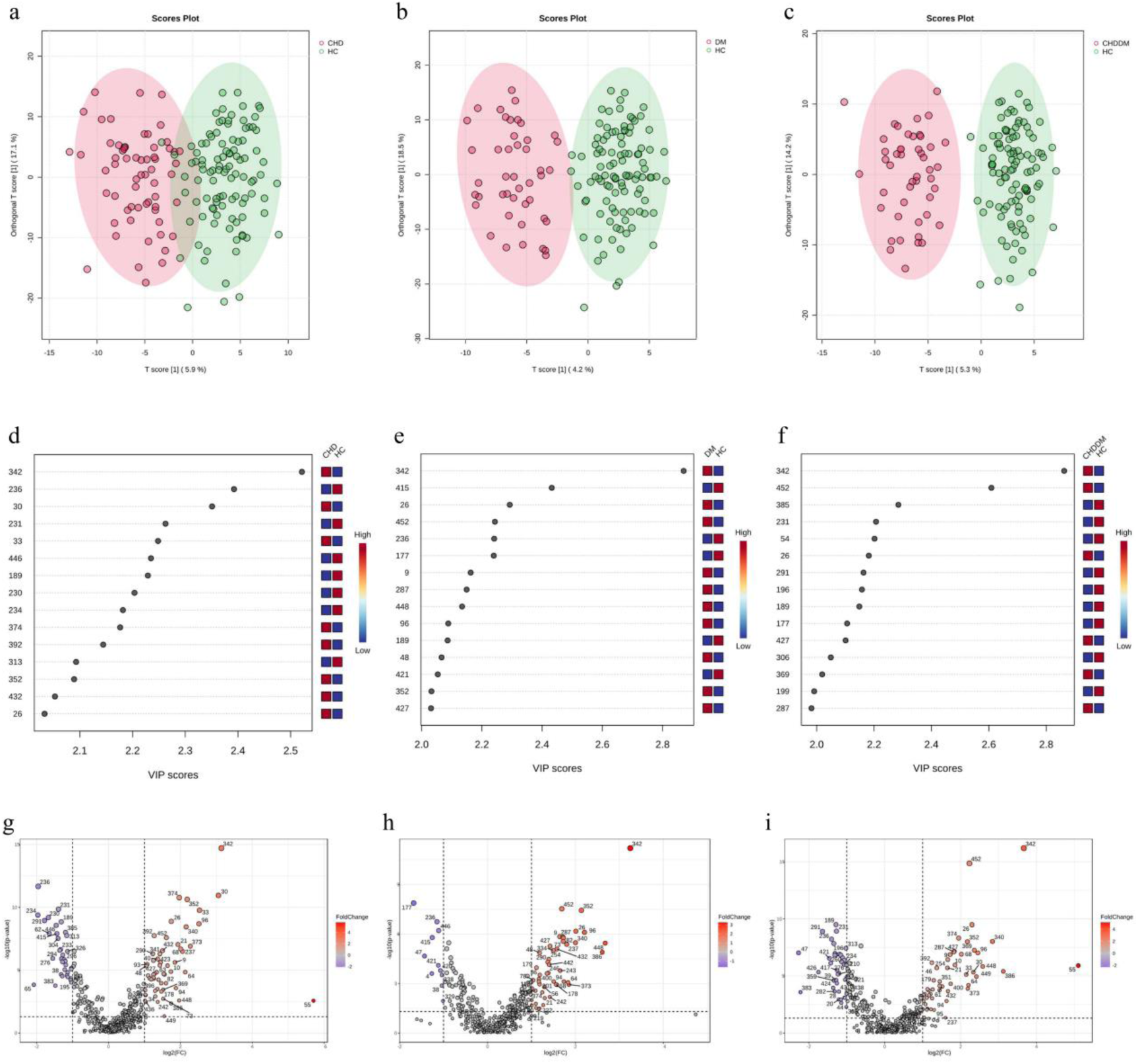
Multivariate analysis for comparison between patients and healthy controls. Orthogonal partial least-squares discriminant analysis (OPLS-DA) score plots were displayed with 95% confidence regions for differentiation of HC and patients with CHD (a), DM (b), and CHDDM (c), respectively. Green, HC samples; Red, disease samples. OPLS-DA-based variable importance in projection (VIP) analysis for the metabolic features contributing to discrimination of patients with CHD (d), DM (e), and CHDDM (f) from HC, respectively. VIP score indicates relative contribution of the metabolites. The top 15 high contribution metabolites are presented. The significantly changed metabolites were displayed with *p*-value < 0.05 and |Log_2_(FC)| > 1 in patients with CHD (g), DM (h), and CHDDM (i) compared to HC, respectively. Red indicates an increase, while blue indicates a decrease. Metabolites were represented in number (see Table S1).

### Identification of potential biomarker for the diseases

Building on the successful distinction between disease groups and HC, we aimed to identify specific metabolites contributing to the differentiation. Variable importance in projection (VIP)-plots based on the OPLS-DA models (**Figure 1d-f**) and volcano plots (**Figure 1g-i**) were constructed. Using the criteria of *p*-value < 0.05, |log_2_(FC)| > 1, and VIP > 1.5, we identified 63, 27, 56 disease-associated metabolic biomarkers for CHD, DM, and CHDDM, respectively. Among these, 11 metabolites were shared across all diseases (see **Table S3**), including 7 lipids (e.g. ceramide, sphingomyelin, fatty acids, and phosphatidylethanolamine (PE)), 2 carboxylic acids (pyruvic acid and pyroglutamic acid that involved in energy metabolism), and 2 nucleotides (hypoxanthine and xanthine related to purine metabolism).

Additionally, 6 metabolites, such as L-glutamic acid, alpha-ketoglutaric acid, tryptophan, glycerophosphocholine, PE 38:6, and L-prolyl-L-threonine, were common to both DM and CHDDM. Dehydroisoandrosterone-3-sulfate, D-glucosamine-6- phosphate, and 4 lipids (such as SM d40:1 (d18:1/22:0), LysoPC 16:1 (16:1/0), Cer d35:1 (d18:1/17:0), and PI 34:2) were shared between CHD and DM. 10 metabolites, including ornithine, choline, L-histidine, allantoic acid, D-galactonic acid, N-acetyl-L-methionine, N-acetyl-putrescine, N-formyl-L-methionine, L-leucyl-L-leucine, L-valyl-L-alanine, and 11 lipids (e.g. phosphatidylcholine (PC), PE, and sphingomyelin) were common to CHD and CHDDM.

Moreover, 1-methylnicotinamide, glycochenodeoxycholic acid, and two types of long-chain fatty acids (such as palmitoleic acid and heptadecenoic acid (17:1)) were unique to DM. Arginine, (R)-2-aminobutanoic acid, fructose, L-phenylalanyl-L-phenylalanine, and a total of 14 different types of lipids were unique to CHDDM. Gluconic acid lactone, 5’-methylthioadenosine, L-kynurenine, acetyl-L-lysine, pyrrole-2-carboxylic acid, and a total of 20 lipids were unique to CHD. These results highlighted that the distinct metabolic changes were associated with each specific disease.

### Disease-related metabolic pathway

To evaluate disrupted metabolic pathways, we selected hydrophilic metabolites with HMDB IDs and conducted a pathway enrichment analysis. A comprehensive examination of the disrupted metabolic pathways identified several significantly altered pathways (p < 0.05) shared across all diseases (**Figure 2**), including arginine and proline metabolism, alanine, aspartate, and glutamate metabolism, and glutathione metabolism. Purine metabolism and glycine, serine, and threonine metabolism were enriched in both the CHD and CHDDM groups. Arginine biosynthesis, butanoate metabolism, citrate cycle (TCA cycle), lipoic acid metabolism, and glyoxylate and dicarboxylate metabolism were disturbed in the DM and CHDDM groups. Interestingly, beside the pathways shared by all 3 diseases, no additional enriched pathways were common between CHD and DM groups. Additionally, unique pathways were identified for each condition: nitrogen metabolism for DM, cysteine and methionine metabolism for CHD, and histidine metabolism and glycerophospholipid metabolism for CHDDM. These findings suggest that while some pathways are commonly disrupted across these diseases, others are more specific to individual conditions.

**Figure 2.**
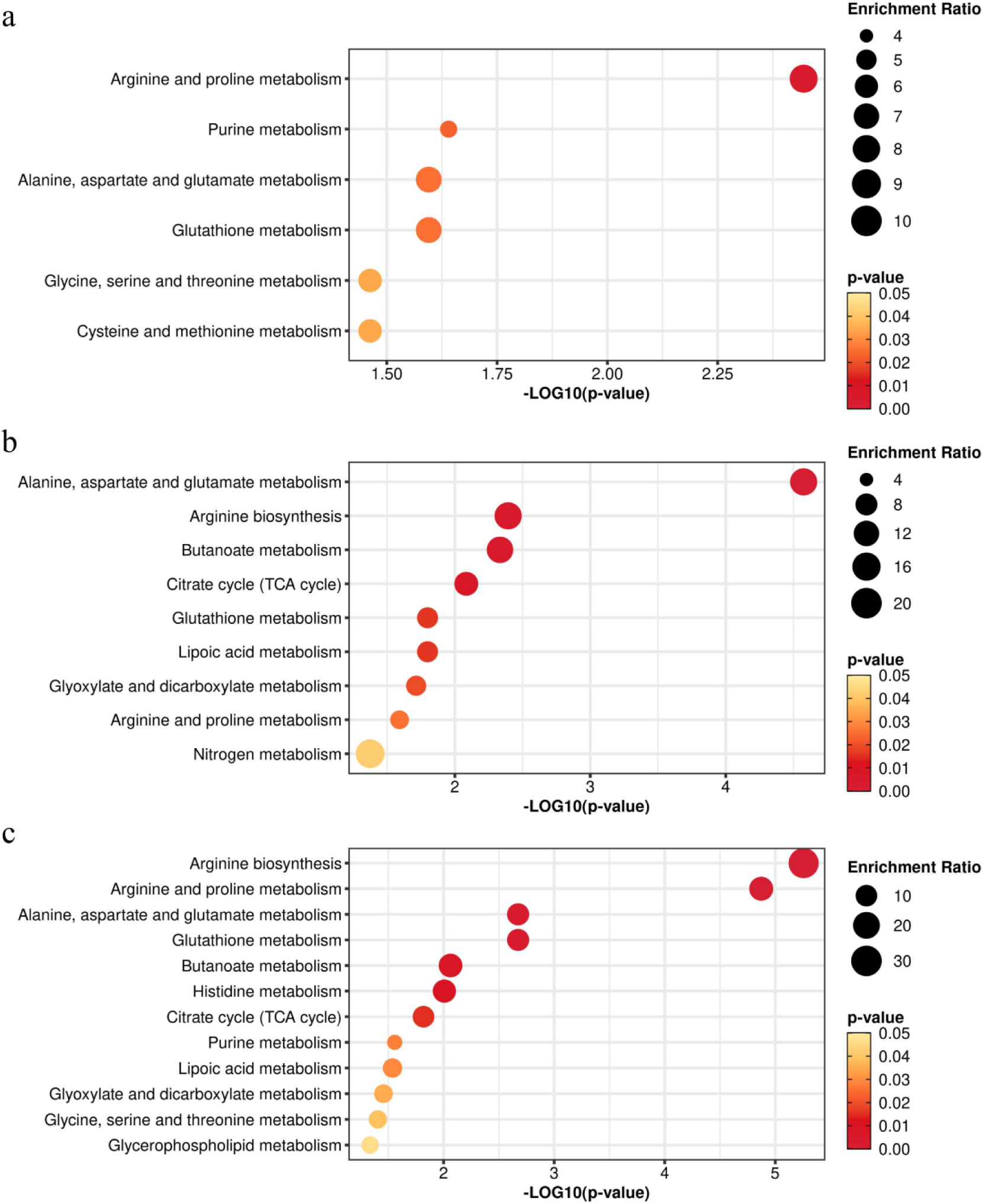
The disturbed metabolic pathways. Pathway enrichment analysis based on differential metabolites between HC and patients with CHD (a), DM (b), and CHDDM (c), respectively. The *p*-value of showed pathways is less than 0.05. Six, nine, and twelve perturbed metabolic pathways were indicated for patients with CHD, DM, and CHDDM, respectively.

### Specific biomarkers for disease diagnosis

To assess the diagnostic potential of dysregulated metabolites, we screened 63, 27, and 56 disease-associated metabolic biomarkers using the area under the curve (AUC) from univariate ROC analysis. Applying an AUC cutoff value > 0.77, we identified 10, 4, and 8 disease-specific biomarkers for CHD, DM, and CHDDM, respectively (see **Table S1** and **Figure 3a** and **3b**). Notably, hypoxanthine was a shared biomarker across all three diseases, with AUC values of 0.848 (95% CI, 0.788–0.906) for CHD, 0.852 (95% CI, 0.786–0.921) for DM, and 0.777 (95% CI, 0.7–0.85) for CHDDM. Other metabolites, pyruvic acid was common to DM and CHDDM, with AUC values of 0.871 (95% CI, 0.805–0.931) for CHDDM and 0.777 (95% CI, 0.685–0.853) for DM. Choline, pyroglutamic acid, PE 36:3p (18:1p/18:2), and PC 34:2e (16:0e/18:2) were shared between CHD and CHDDM, exhibiting AUC values greater than 0.78 in both CHDDM and CHD, but as low as 0.677 in the DM group. Among other findings, glycerophosphocholine and turanose were distinctive to DM, xanthine and PE 38:5p (18:1p/20:4) were aberrant for CHDDM, and N-Formyl-L-methionine, N-Acetyl-L-methionine, PE 36:1p (18:0p/18:1), PE 36:2p (18:0p/18:2), and PE 34:2p (16:0p/18:2) were predominant in CHD. These disease-unique metabolites demonstrated superior diagnostic performance in their respective conditions compared to their performance in other diseases (**Figure 3b**). Consequently, we conducted multivariate ROC analysis using a panel of unique biomarkers, expecting that the disease-unique markers would exhibit a stronger synergistic response within their corresponding diseases (**Figure 3c-e**). The results revealed that the metabolic panels of these unique biomarkers achieved an AUC of 0.935 with 0.938 sensitivity at 0.8 specificity in CHD, an AUC of 0.913 with 0.909 sensitivity at 0.72 specificity in DM, and an AUC of 0.903 with 0.833 sensitivity at 0.84 specificity in CHDDM (see **Table S4**) in test set, demonstrating higher performance in their respective diseases compared to others. The results indicated that these unique metabolite panels are high accuracy for diagnosing CHD, DM, and CHDDM.

**Figure 3.**
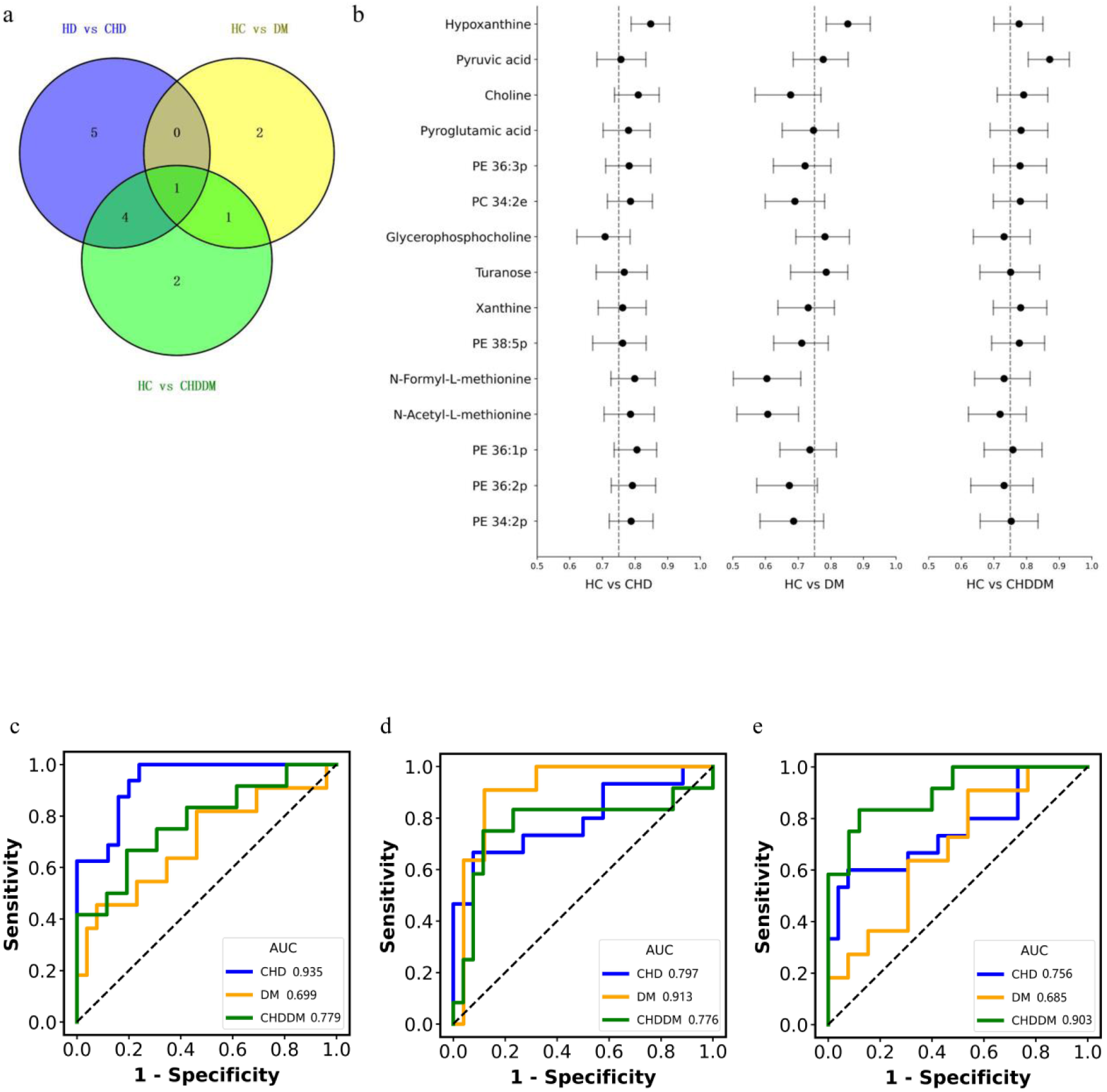
Identification of potential disease-specific biomarkers. (a) Venn diagram presents differential metabolites in patients with CHD, DM, and CHDDM compared to HC, respectively. The differential metabolites were selected with *p*-value < 0.05, |Log_2_(FC)| > 1, VIP > 1.5, and AUC values of univariate ROC > 0.77 in patients with CHD, DM, and CHDDM, compared to HC, respectively. (b) Univariate ROC analysis was performed based on the machine-learning algorithm of support vector machines (SVM) in patients with CHD, DM, and CHDDM, compared to HC, respectively. Data are presented as AUC at 95% confidence intervals (CIs) in training set. The ROC was established in test set to examine the performance of the unique metabolite biomarker panels obtained from CHD (c), DM (d), and CHDDM in distinguishing patients with CHD (blue), DM (yellow), and CHDDM (green), compared to HC, respectively.

### Metabolic biomarkers for assessing CHD risk in DM patients

Given the increased risk of CHD in DM patients, we sought to identify metabolic markers that could predict this risk. By focusing on metabolites significantly altered between DM and CHDDM and excluding those primarily affected by CHD, we identified 23 potential biomarkers (**Figure 4a, Table S5**). Multivariate ROC analysis showed that these biomarkers effectively distinguished between DM and CHDDM, with AUC values of 0.864 (95% CI, 0.639–1.0) in the training set and 0.862 (95% CI, 0.676–0.984) in the test set (**Figure 4b**). In contrast, the same markers yielded AUC values of only 0.547 (95% CI, 0.333–0.763) and 0.518 (95% CI, 0.326–0.701) in the training and test sets between HC and CHD (**Figure 4c**). These findings suggested that these 23 metabolites could serve as early indicators of CHD risk in DM patients.

**Figure 4.**
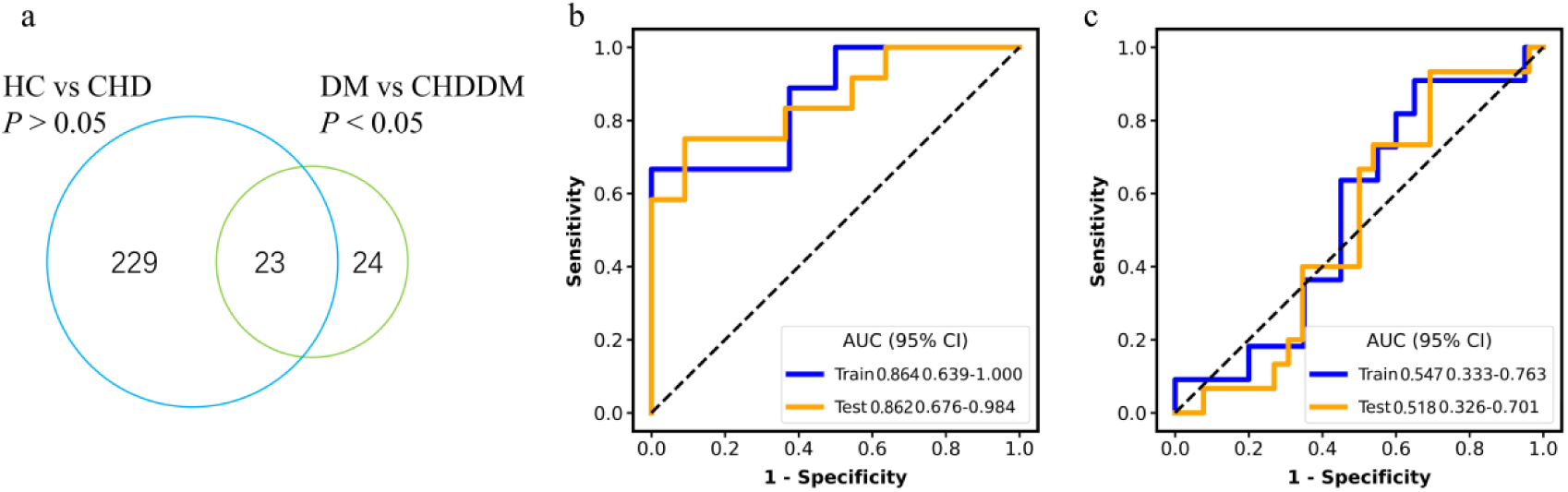
The metabolic biomarkers for patients with DM susceptible of CHD. (a) Venn diagram of metabolites from significantly altered in CHDDM compared to DM and insignificant biomarkers between HC and CHD, respectively. (b) Multivariate ROC analysis was performed in training set (blue) and test set (yellow) based on the machine-learning algorithm of support vector machines (SVM) in DM vs CHDDM (b) and HC vs CHD (c) patients group, respectively. Data are presented as AUC at 95% confidence intervals (CIs).

## Discussion

This study employed a multi-platform metabolomic approach to profile plasma metabolites across CHD, DM, and CHDDM. We identified 63, 27, and 56 disease-associated metabolic biomarkers for CHD, DM, and CHDDM, respectively, and performed pathway enrichment analysis to uncover disrupted metabolic pathways. Several unique biomarkers demonstrated strong diagnostic potential, with 23 metabolites identified as potential indicators of CHD risk in DM patients.

One of the key findings was the accumulation of hypoxanthine across all diseases. Hypoxanthine, a purine derivative, is a product of adenosine triphosphate degradation and is eventually excreted as uric acid (19). Elevated uric acid is strongly associated with the development of diabetes (20). In both CHD and DM, elevated oxidative stress leads to increased hypoxanthine levels due to enhanced purine metabolism (21). This oxidative stress contributes to endothelial dysfunction and inflammation, exacerbating cardiovascular and metabolic diseases(22).While hypoxanthine is elevated in both conditions, it is more closely linked to ischemia-reperfusion injury in CHD and reflects chronic hyperglycemia and insulin resistance in DM (23). In our study, uric acid and hypoxanthine were significantly increased across CHD, DM, and CHDDM. Hypoxanthine has been identified as a significant differential metabolite in CHDDM compared to CHD (9). Although we did not perform a direct comparison between CHD and CHDDM, the fold change analysis indicates that hypoxanthine levels in CHDDM are 1.34 times higher than in CHD, emphasizing the critical role of oxidative stress and purine metabolism in the development and progression of these conditions.

Pyruvic acid, is a crucial intermediate in glucose metabolism, also emerged as a common metabolic marker in DM and CHDDM, indicating disruptions in glycolytic flux and mitochondrial function. Elevated plasma pyruvic acid levels have been associated with increased insulin resistance in DM patients (24). In CHDDM, pyruvic acid accumulation is linked to heightened oxidative stress and inflammation, key factors in the pathogenesis of both CHD and DM. Elevated pyruvic acid levels in CHDDM patients correlate with poorer cardiovascular outcomes and may indicate greater disease severity (25).

Choline, another identified metabolite, acts as a precursor to trimethylamine N-oxide (TMAO), a metabolite that has emerged as a potential biomarker for cardiovascular diseases(26). Elevated levels of TMAO are associated with an increased risk of adverse cardiovascular events, primarily due to its role in promoting atherosclerosis and thrombosis. These findings underscore the significance of choline metabolism—and consequently TMAO—in the understanding and management of CHD and CHDDM (27). Pyroglutamic acid, also known as 5-oxoproline, involved in the glutathione cycle and associated with oxidative stress, was also identified. Elevated levels of pyroglutamic acid may indicate impaired glutathione metabolism, a significant factor in oxidative stress, a common pathway contributing to both CHD and CHDDM (28).

Glycerophospholipids, such as PE 36:3p (18:1p/18:2) and PC 34:2e (16:0e/18:2), are critical for cellular membranes. Specific glycerophospholipid species, including phosphatidylcholines (PCs) and phosphatidylethanolamines (PEs), are significantly altered in patients with CHD (29) and are associated with the severity of atherosclerosis, myocardial infarction (MI), and overall cardiovascular risk (30).

The metabolic biomarkers shared among these diseases reflect the commonality between them, highlighting their association with metabolic syndrome. However, when it comes to distinguishing and diagnosing these diseases, the aforementioned metabolites might exhibit lower efficiency (see **Figure 3b**). Therefore, unique metabolic markers of each disease are more effective in distinguishing and diagnosing the disease accurately. The present study identified several disease-specific metabolites, for instance, glycerophosphocholine and turanose were distinctive to DM, xanthine and PE 38:5p (18:1p/20:4) were aberrant for CHDDM, and N-Formyl-L-methionine, N-Acetyl-L-methionine, PE 36:1p (18:0p/18:1), PE 36:2p (18:0p/18:2), and PE 34:2p (16:0p/18:2) were predominant in CHD (**Figure 3a and** **3b**).

Glycerophosphocholine (GPC) has been identified as a potential biomarker for insulin resistance, a precursor to type 2 diabetes (T2DM). Studies have shown that GPC levels are inversely related to insulin sensitivity. In particular, GPC, along with other metabolites, was found to be associated with the risk of developing glucose intolerance, suggesting its role in early detection of metabolic disturbances leading to T2DM (31,32). Turanose, a disaccharide, is recognized for its low glycemic index, making it a potential option for managing blood sugar levels in diabetic patients. Although turanose has not yet been reported as a biomarker for DM diagnosis, its unique metabolic properties make it a safer alternative to common sugars, contributing to improved glycemic control in individuals with or at risk of diabetes (33).

Xanthine, a product in the purine degradation pathway, has been investigated for its role in cardiovascular and metabolic diseases. The enzyme xanthine oxidoreductase (XOR) converts xanthine to uric acid, a process linked with increased oxidative stress. Elevated XOR activity and subsequent xanthine accumulation are associated with various metabolic conditions, including CHD and DM. Studies have demonstrated that increased plasma XOR activity is correlated with higher BMI, elevated liver enzymes, and other markers of metabolic dysfunction, suggesting that xanthine could be a valuable biomarker in CHDDM (34).

PE species, such as PE 38:5p, are essential components of cell membranes and are involved in lipid metabolism and signaling. In cardiovascular disease and diabetes, changes in PE levels have been associated with metabolic syndrome, insulin resistance, and atherosclerosis. The profile of PE 38:5p, including its fatty acid composition, may indicate underlying metabolic disturbances in CHDDM, making it a potential marker for disease progression and risk assessment (35).

N-Formyl-L-methionine and N-Acetyl-L-methionine are derivatives involved in methionine metabolism with significant roles in cellular processes. Elevated levels of N-Formyl-L-methionine and N-Acetyl-L-methionine may reflect disruptions in methionine metabolism, which is important for cardiovascular health(36,37). Phosphatidylethanolamine plasmalogens (PE 36:1p, PE 36:2p, PE 34:2p) are phospholipids crucial for membrane structure and function, particularly in the heart and brain. Alterations in these plasmalogens are associated with oxidative stress and inflammation, key factors in the progression of CHD. Decreased levels of plasmalogens, such as PE 36:1p, PE 36:2p, and PE 34:2p, have been linked to an increased risk of cardiovascular disease due to their role in protecting cells from oxidative damage and maintaining membrane integrity (38).

These metabolic biomarkers not only perform well individually within their respective diseases, but when used together as a panel, they offer superior diagnostic performance within their respective conditions compared to others (**Figure 3c-e**). In practical diagnostic steps within a hospital setting, the initial task is to determine whether a disease is present, followed by classifying it as CHD, DM, or CHDDM. The disease-specific metabolic biomarkers mentioned above can effectively distinguish between these conditions.

However, none of the specific biomarkers identified for CHD, DM, and CHDDM overlapped with those we discovered (39). Moreover, the identified disease-specific metabolic biomarkers in previous study exhibited high efficacy both within their respective conditions and across other disease. This discrepancy likely results from the differing detection methodologies, as the previous study employed nuclear magnetic resonance (NMR), identifying only 42 metabolites—less than one-tenth of the 453 metabolites detected in our study. The broader metabolite coverage in our analysis provides a distinct advantage for biomarker identification and pathway analysis.

The present study also identified several disease-related metabolic pathways (see **Figure 2a-c**). Arginine and proline metabolism, alanine, aspartate, and glutamate metabolism, as well as glutathione metabolism, were enriched across all diseases. Arginine and proline metabolism is crucial for nitric oxide synthesis, which is essential for vascular function. Disruptions in these pathways are associated with endothelial dysfunction, impaired energy metabolism, and increased oxidative stress, contributing to the progression of both cardiovascular and diabetic complications. Several studies have indicated that arginine and proline metabolism is significantly altered in the DM group compared to HC (40) and underwent remarkable changes in CHD group (41), consistent with the observations in our study. Previous studies have shown that alanine, aspartate, and glutamate metabolism are extensively altered in CHD (42) and DM (43), consistent with our findings. These pathways play a crucial role in amino acid metabolism and energy production, and disruptions can lead to impaired energy metabolism, a common characteristic of both CHD and DM. Elevated levels of metabolites from these pathways have been observed in patients with CHD and DM, indicating their involvement in the pathophysiology of these diseases. As a major antioxidant pathway, glutathione metabolism is essential for managing oxidative stress, which is heightened in CHD and DM. Dysregulation in this pathway can lead to increased oxidative damage, contributing to the progression of both cardiovascular and diabetic complications (44,45).

Additionally, several unique metabolic pathways were identified: nitrogen metabolism for DM, cysteine and methionine metabolism for CHD, and histidine metabolism and glycerophospholipid metabolism for CHDDM. Previous studies have shown that maintaining nitrogen balance can prevent the progression of diabetes (46), and reduced nitrogen synthesis has been associated with diabetes (47). These findings suggest that nitrogen metabolism may play a critical role in the progression of diabetes and its complications.

Cysteine and methionine metabolism play significant roles in cardiovascular health, particularly in the context of CHD. These pathways are involved in the production of key molecules such as glutathione, a major antioxidant that helps protect cells from oxidative stress, which is a known contributor to CHD (41). Disruptions in cysteine and methionine metabolism can lead to an imbalance in redox status, elevated homocysteine levels, and increased oxidative stress, all of which are associated with a higher risk of atherosclerosis and other cardiovascular complications (48). Cysteine protease inhibitor C can be considered as another predictor of cardiovascular complications, indirectly indicating that cysteine metabolism plays an important role in the pathogenesis and prognosis of coronary heart disease (49).

Histidine, an essential amino acid, is involved in several critical metabolic processes, including the synthesis of histamine, which plays a key role in immune response, gastric acid secretion, and neurotransmission (50). Studies have shown that histidine is associated with the risk of developing T2D and CHD (51). In the context of CHD and DM, dysregulation of histidine metabolism may contribute to the chronic inflammatory state commonly observed in these patients, potentially worsening cardiovascular complications (52).

For DM patients at risk of CHD, certain metabolic biomarkers can serve as early indicators of increased cardiovascular risk. By focusing on metabolites significantly altered between DM and CHDDM, while excluding those primarily affected by CHD, we isolate metabolic changes specifically linked to the progression from DM to CHDDM, improving the specificity of biomarkers for early detection and targeted intervention. To our knowledge, this is the first study to report 23 metabolites that could serve as potential biomarkers for assessing the risk of CHD in DM patients. These findings highlight the importance of metabolic profiling for understanding disease mechanisms and developing precise diagnostic tools.

### Study limitations

First, this study is a single-center study, and considering its suitability for the general population, the scope may be expanded to multi-center and other species in the future. Second, this study focuses on plasma metabolomics in patients with CHD, DM, and CHDDM. The combination of multiple omics data, such as genomics and proteomics, can provide stronger insights, which can better elucidate the disease mechanism, promote people’s in-depth understanding and better characterization of CHD, DM, and CHDDM. Third, current research is focused on characterizing the disease status of patients, and further basic medical validation is needed to elucidate the mechanism of CHD, DM, and CHDDM. In future studies, further validate the metabolic markers we screened through basic experiments such as animal experiments and cell experiments, etc, ultimately better serving clinical work.

## Conclusion

This study advances our understanding of the metabolic underpinnings of CHD and DM, particularly when these conditions coexist as CHDDM. Through multi-platform metabolomics, we identified specific biomarkers unique to each condition, with several demonstrating strong diagnostic potential. By isolating metabolites associated with the progression from DM to CHDDM, we identified 23 potential biomarkers for assessing CHD risk in DM patients. These findings underscore the value of metabolic profiling in developing precise diagnostic tools and therapeutic strategies for CHD, DM, and CHDDM.

## Acknowledgements

The authors thank all the patients, volunteers, and their families, the investigators, and the site personnel who participated in this study.

## Sources of Funding

This work was supported by grants from the National Natural Science Foundation of China (82070329 and 82170456).

## Declaration of interests

Fei Shi, Hui-Na Cui, Yun He, Li Liu, Yan Li, and He Wen are employees of Metanotitia Inc., Harbin. Zhicai Wang, Yan Cang, Yi Zhang, Yawei Xu, Zheng Liu have nothing to declare.

## Conflict of Interests

The authors declare that there is no conflict of interest regarding the publication of this paper.

## Ethics approval and consent to participate

This study was approved by the Research Ethics Committee of the Tenth People’s Hospital of Shanghai, No. 20K149 and 21K65, and the informed consents were obtained from all participants. The study was conducted in accordance with the guidelines of the International Conference on Harmonization for Good Clinical Practice and the Declaration of Helsinki.

## Consent for publication

All authors have agreed to publish this manuscript.

## Author contributions

**Conceptualization**: LL, YL, HW, and ZL; **Methodology**: ZW, YC, YH, and YX; **Investigation**: FS and HW; **Resources**: ZW, YC, and YZ; **Visualization**: FS and HC; **Funding acquisition**: ZL; **Writing - Original Draft**: HW and ZL; **Supervision**: YL and ZL. All authors were involved in the review and final approval of the manuscript.

## Data availability

The datasets used and/or analysed during the current study available from the corresponding author on reasonable request.

## Abbreviation

CHD: coronary heart disease
DM: diabetes mellitus
HC: healthy controls
BMI: body mass index
BUN: blood urea nitrogen
LDL: low-density lipoprotein
HDL: high-density lipoprotein
BMI: body mass index
TG: triglycerides
HbA1c: glycated hemoglobin
CV: coefficient of variation
PC: phosphatidylcholine
PE: phosphatidylethanolamine
LysoPC: lysophosphatidylcholine
LysoPE: lysophosphatidylethanolamine
MePC: monoether phosphatidylcholine
AcCa: Acetyl-CoA Carboxyltransferase α
Cer: ceramide
PI: phosphatidylinositol
PS: Phosphatidylserine
SM: Sphingomyelin
ChE: Cholesterols ester
FA: fatty acid
DAG: diacylglycerol
TAG: triacylglycerol
GC-MS: gas chromatography-mass spectrometry
UHPLC-MS: ultra-high performance liquid chromatography-mass spectrometry
QC: quality control
OPLSDA: orthogonal projections to latent structures discriminant analysis
VIP: variable importance in projection
ROC: receiver operating characteristic
AUC: area under the ROC curve
BCAAs: branched-chain amino acids
TOF: time-of-flight
ESI: electrospray ionization
DDA: data-dependent acquisition
FAME: fatty acid methyl esters
RT: retention time
NormAE: normalization autoencoder
SVM: support vector machines
CIs: confidence intervals

## Notes

### Competing Interest Statement

The authors have declared no competing interest.

